# Linear causal filtering: definition and theory

**DOI:** 10.1101/2021.05.01.442232

**Authors:** Roberto D. Pascual-Marqui, Rolando J. Biscay, Jorge Bosch-Bayard, Peter Achermann, Pascal Faber, Toshihiko Kinoshita, Kieko Kochi

**Affiliations:** The KEY Institute for Brain-Mind Research, Department of Psychiatry, Psychotherapy, and Psychosomatics, University Hospital of Psychiatry, Zurich, Switzerland; Centro de Investigacion en Matematicas, Guanajuato, Mexico; Montreal Neurological Institute, McGill University, Montreal, Canada; Department of Neuropsychiatry, Kansai Medical University, Osaka, Japan

## Abstract

This work provides a framework based on multivariate autoregressive modeling for linear causal filtering in the sense of Granger. In its bivariate form, the linear causal filter defined here takes as input signals A and B, and it filters out the causal effect of B on A, thus yielding two new signals only containing the Granger-causal effect of A on B. In its general multivariate form for more than two signals, the effect of all indirect causal connections between A and B, mediated by all other signals, are accounted for, partialled out, and filtered out also. The importance of this filter is that it enables the estimation of directional measures of causal information flow from any non-causal, non-directional measure of association. For instance, based on the classic coherence, a directional measure of strength of information flow from A to B is obtained when applied to the linear causal filtered pair containing only A to B connectivity information. This particular case is equivalent to the isolated effective coherence (doi.org/10.3389/fnhum.2014.00448). Of more recent interest are the large family of phase-phase, phase-amplitude, and amplitude-amplitude cross-frequency coupling measures which are non-directional. The linear causal filter makes it now possible to estimate the directional causal versions these measures of association. One important field of application is in brain connectivity analysis based on cortical signals of electric neuronal activity (e.g. estimated sources of EEG and MEG, and invasive intracranial ECoG recordings). The linear causal filter introduced here provides a novel solution to the problem of estimating the direction of information flow from any non-directional measure of association. This work provides definitions, non-ambiguous equations, and clear prescriptions for implementing the linear causal filter in diverse settings.

## 2. The multivariate autoregressive model

It is assumed throughout that the set of time series recordings have no measurement noise, and that there are no other sources of influence other than those expressed by the multivariate autoregressive model. Furthermore, the time series are assumed to be wide sense stationary, with zero mean. Informally, this means that the multivariate autoregressive model fully describes the measurements, and that it can be estimated given sufficient data.

The methods described here apply to zero mean signals that are sampled in the form of *N_E_* epochs, each consisting of *N_T_* discrete time samples, i.e. *t* = 0…(*N_T_* − 1).

For the multivariate time series **x***(t*) ∈ ℝ^*p*×1^, for *t* = 0…(*N_T_* − 1), consider the autoregressive model of order *“q”*:

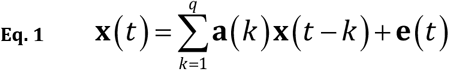

where **e***(t*) ∈ ℝ^*p*×1^ is the innovations time series with covariance matrix **S**_*ee*_ ∈ ℝ^*p*×*p*^, and **a**(*k*) ∈ ℝ^*p*×*p*^ are the autoregressive coefficients for *k* = 1…*q* . It is assumed that *q* ≥ 1 and *p* ≥ 2 . Furthermore, there should be sufficient data for estimation.

Given time series recordings for **x**(*t*) and the autoregressive order “*q*”, estimators for the coefficients **a**(*k*) in Eq. 1 can be obtained by, e.g., least squares (see e.g. Shumway and Stoffer 2017).

The frequency domain representation of Eq. 1 (see e.g. Wei 2019) is:

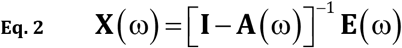

where **I** is the identity matrix of appropriate dimension (in this case **I** ∈ ℝ^*p*×*p*^), **X**(ω) ∈ ℂ^*p*×1^ and **E**(ω) ∈ ℂ^*p*×1^ are the discrete Fourier transforms of **x***(t*) and **e***(t*) respectively, ω denotes the discrete frequency, and **A**(ω) ∈ ℂ^*p*×*p*^ is the discrete Fourier transform of the extended sequence **a**(*k*), defined as zero for *k* = 0 and for *k* > *q*, i.e.:

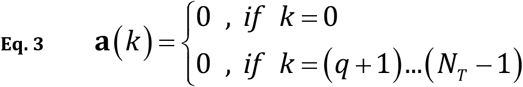

For the sake of completeness, the discrete Fourier transform is defined in Eq. 60 in “Appendix A”. A simplified equation for the sequence of autoregressive coefficients is presented in Eq. 62 in “Appendix A”.

The cross-spectrum is (see e.g. Wei 2019):

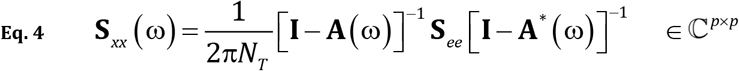

where the superscript “*” denotes transposed and complex conjugate.

Note that the Fourier transform of the innovations can be obtained from Eq. 2:

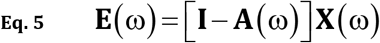

## 3. Causality in the sense of Granger

Throughout, the term “causal” is exclusively limited to causality in the sense of Granger. Causality in the sense of Granger is defined by non-zero coupling coefficients. For instance, if:

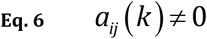

for some pair of signals (*i* ≠ *j*), for some lag *“k”,* then *“j* Granger causes *i”.* In Eq. 6, *a_ij_* (*k*) is the corresponding element of **a**(*k*). This translates to the frequency domain, where “*j* Granger causes *i*” if:

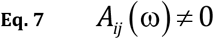

for some pair of signals (*i* ≠ *j*), for some frequency “ω”. In Eq. 7, *A_ij_* (ω) is the corresponding element of **A**(ω).

## 4. Least squares estimation for the multivariate autoregressive model: coefficients and innovations covariance

The multivariate autoregressive model of order *“q”* for zero mean time series recordings **X**_ε_(*t*) ∈ ℝ^*p*×1^, sampled in the form of *N_E_* epochs, each consisting of *N_T_* discrete time samples is:

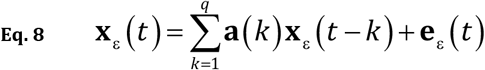

for ε = 1…*N_E_* and *t* = *q*…(*N_T_* − 1). This can be written as:

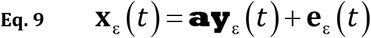

with:

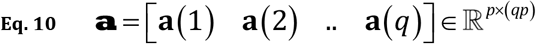

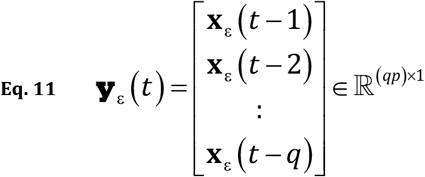

The least squares solution is:

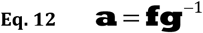

with:

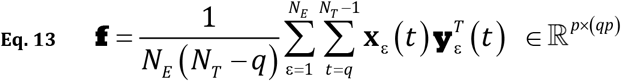

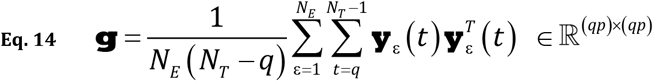

The estimator for the covariance matrix of the innovations **e**(*t*) in Eq. 8 and Eq. 9 is:

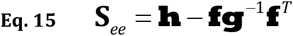

where:

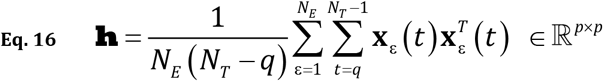

Note that these equations are valid for stable autoregressive processes if the inverse of **g** in Eq. 14 exists.

As usual, the order “*q*” can be estimated using Akaike’s information criterion (AIC) (Akaike 1974).

## 5. The linear causal filter

The linear causal filter defined here takes as input a multivariate time series, and for any given distinct pair of signals *“i”* and *“j”,* it filters out the effect of all causal connections mediated by all other signals (accounting for, and partialling out their effects), and it filters out the causal effect of “*j*” on “*i*”, thus leaving exclusively the effect of “*i*” on “*j*”.

The derivation of the linear causal filter is presented in three steps.

In a first step, the discrete Fourier transform of the multivariate time series, given by Eq. 2, is reduced to the two variables of interest, namely the *i*-th and the *j*-th time series, giving:

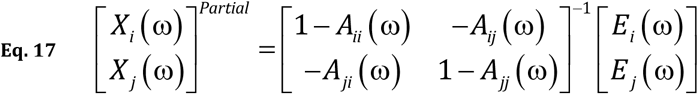

for any *i* ≠ *j* with *i*, *j* = 1…*p*, where *A*_●●_ (ω), *E*_●_ (ω), and *X*_●_ (ω) correspond to the elements of **A**(ω) ∈ ℂ^*p*×*p*^, **E**(ω) ∈ ℂ^*p*×*p*^, and **X**(ω) ∈ ℂ^*p*×1^ respectively.

In the first step above, all indirect causal connections between the *i*-th and *j*-th variables have been accounted for and partialled out, yielding the simple system in Eq. 17. This is true because the coefficients and innovations in the right-hand side of Eq. 17 are multivariate, and not marginal bivariate (except for the case when the total number of signals is exactly two).

In the second step, the directional connection from “*j* to *i*” is severed (i.e. cut off), leaving exclusively the direct, directional connection from *“i* to *j*”, giving:

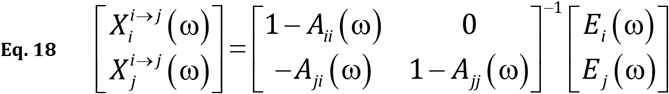

in which the coupling coefficient from “*j* to *i*” has been set to zero, i.e.:

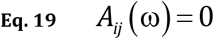

while all other coefficients and innovations remain unchanged.

It is of interest to note that the coherence computed from Eq. 18 is identical to the isolated effective coherence for “*i* to *j*” in (Pascual-Marqui et al 2014) for independent innovations.

Eq. 18 still does not have the form of a general filter, which takes as input the multivariate data **X**(ω) ∈ ℂ^*p*×1^ and returns as output the bivariate system 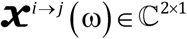 containing exclusively information flow from *“i* to *j*”.

In the third step, linear causal filtering is expressed as:

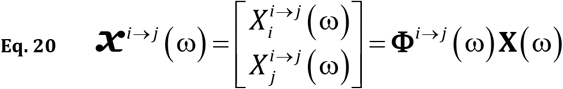

by plugging Eq. 5 into Eq. 18.

In Eq. 20, the linear causal filter **Φ**^*i*→*j*^ (ω) ∈ ℂ^2×*p*^ is:

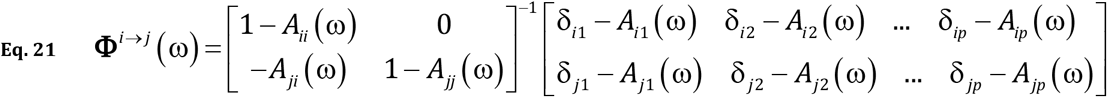

where δ_*ij*_ is the Kronecker delta, defined as:

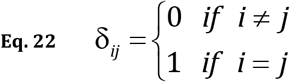

The linear causal filter is completely determined by the multivariate autoregressive coefficients, as can be seen from its definition in Eq. 21.

Linear causal filtering with Eq. 20 will give the discrete Fourier transforms 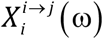 and 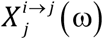 containing only direct, directional information flow from *“i* to *j”*. The inverse discrete Fourier transforms of 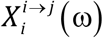 and 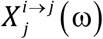 will produce the real-valued filtered time series, denoted 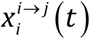 and 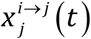, which contains only direct, directional information flow from “*i* to *j*”. For the sake of completeness, “Appendix B” defines the inverse discrete Fourier transform in general and includes information on how to use it when the output is a real signal with DC level equal to zero, which is the case of interest here.

Note that by definition, in the bivariate case (i.e. only two signals total), the linear causal filter for “*i*(=1) to *j*(=2)” will return new signals that have zero coupling from “*j*(=2) to *i*(=1)”, i.e.:

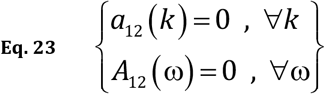

Equations for information flow in the reverse direction from *“j* to *i*” can be written as:

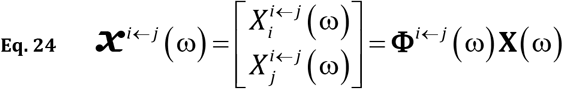

with:

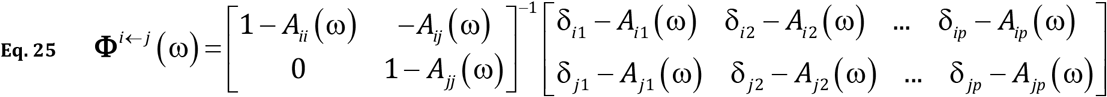

## 6. Generic pipeline for the linear causal filtering of signals

Assumptions about the time series data:

1. Zero mean.
2. No measurement noise.
3. Wide sense-stationarity.
3. There are no other sources of influence other than those expressed by the multivariate autoregressive model.

The following generic algorithm is intended to be simple and straightforward; it is not optimized for efficiency. The algorithm takes in time series data and the autoregressive order, and outputs, for each distinct pair, the filtered signals containing only direct, directional information flow from one to the other, in both frequency and time domains.

**Algl-Step# 1**

Given zero mean multivariate time series data corresponding to electrophysiological signals, **X**_ε_(*t*) ∈ ℝ^*p*×1^, with *p*≥2, for time samples *t* = 0…(*N_T_* − 1) and epochs ε *=* 1…*N*․ Given the autoregressive order *“q”.*

**Alg1-Step# 2**

Estimate the autoregressive coefficients **a**(*k*) ∈ ℝ^*p*×*p*^ for *k* = 1…*q*, using e.g. the least squares method (Eq. 8 through Eq. 14). Noting Eq. 3 above, compute its discrete Fourier transform **A**(ω) ∈ ℂ^*p*×*p*^ (see e.g. “Appendix A”).

**Alg1-Step# 3**

Compute the discrete Fourier transform **X**_ε_ (ω) ∈ ℂ^*p*×1^.

**Algl-Step# 4**

For all *i* = 1…*p* and all *j* = 1…*p*, and if *i* ≠ *j*, do steps #4.1., #4.2., and #4.3:

**Alg1-Step# 4.1.** Compute the causal filter in Eq. 21 and Eq. 22.
**Alg1-Step# 4.2.** Apply the linear causal filter in Eq. 20 to obtain 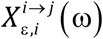 and 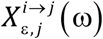 with direct, directional information flow *i* → *j*, for ε = 1…*N_E_*, for ω = 1… (*N_T_*/2).
**Algl-Step# 4.3.** Compute the inverse discrete Fourier transforms (see “Appendix B”) of 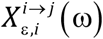 and 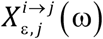 to obtain the real-valued time series with direct, directional information flow 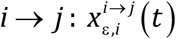 and 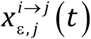.

**Algl-Step# 5**

End.

In summary, this algorithm “Alg1, Steps #1 to #5” takes as input the multivariate time series **X**_ε_(*t*) ∈ ℝ^*p*×1^, and returns for all distinct pairs “*i*” and “*j*”, the following filtered variables where only *i* → *j* information flow remains:

1. The Fourier transforms:

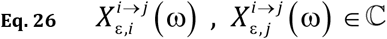
2. The time series

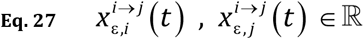

## 7. Instantaneous phase and amplitude (envelope) for linear causal filtered signals

Once the linear causal filtered time series pairs in Eq. 27 are available, they can be further processed by means of the following prescription.

**Alg2-Step# 1**

Band pass filtering of 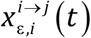 and 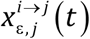 to a frequency band Ω, giving:

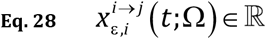

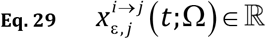

where, e.g. Ω= 5 Hz to 7 Hz for the electrophysiological theta band, or Ω= 35 Hz to 55 Hz for the gamma band (other definitions can be used).

*Note that Ω-band-pass filtering is a post-processing step, it is applied after the linear causal filter, it is not to be applied before the estimation of the multivariate autoregressive model, nor is it to be applied before linear causal filter.*

**Alg2-Step# 2**

Computing the corresponding complex-valued analytic signals (see “Appendix C”), denoted as:

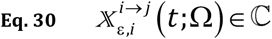

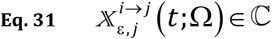

**Alg2-Step# 3**

Computing the envelopes (instantaneous amplitudes):

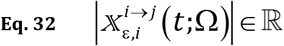

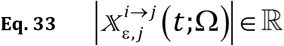

**Alg2-Step# 4**

Computing the instantaneous phases:

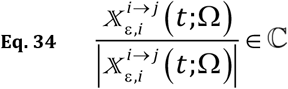

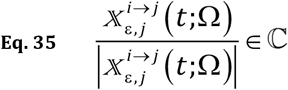

**Alg2-Step# 5**

End.

For the sake of completeness, “Appendix C” contains the definition the Hilbert transform, the analytic signal, the instantaneous phase, and the envelope. Furthermore, of practical value, it contains the algorithm of Marple 1999 for the calculation of the discrete Hilbert transform and of the analytic signal for sampled time series.

## 8. Measures of directional information flow using linear causal filtering

### Directional coherence using linear causal filtering

The squared modulus of the periodogram-based coherence, using the directional Fourier transforms in Eq. 26, is:

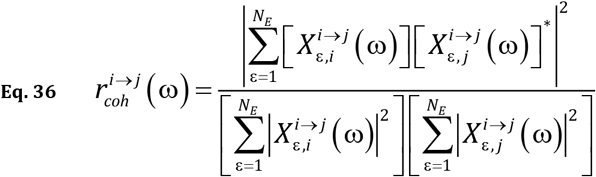

see e.g. Brillinger 2001. This is a directional version of the coherence.

An alternative parametric estimator for directional coherence can be obtained, using the closed form of the cross-spectrum of an autoregressive model. From Eq. 20, the cross-spectral matrix can also be written as:

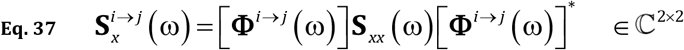

Plugging Eq. 4 into Eq. 37 simplifies to:

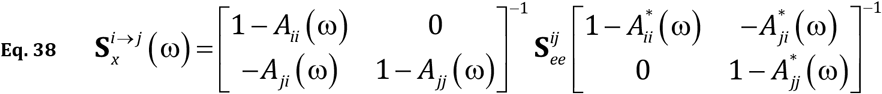

with:

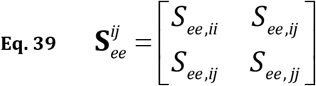

corresponding to the innovations covariance matrix for the *i*-th and *j-th* signals. If the innovations are independent, i.e.:

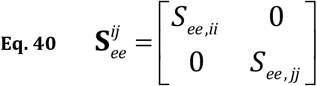

then the squared of the modulus of the coherence giving the directional, causal information flow *“i* to *j*” is:

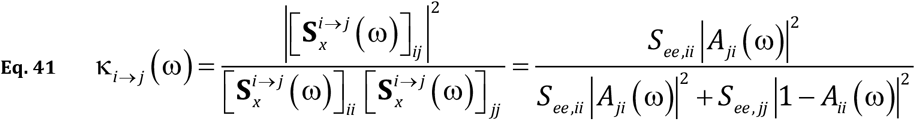

which is the same as the isolated effective coherence (Pascual-Marqui et al 2014).

This measure of directional, causal information flow was shown to have improved properties when compared to other similar measures of directed coherence (see Pascual-Marqui et al 2014). Also included in (Pascual-Marqui et al 2014) are several results illustrating the properties of the isolated effective coherence, based on simulated data and on real human signals of estimated cortical activity.

### Directional phase-phase coupling using linear causal filtering

The non-directional measure of phase-phase coupling between two signals is also known as “phase locking value (PLV)”, “phase synchronization” and “phase coherence”.

Some basic early applied literature on this non-causal, non-directional measure of association are the papers by Tass et al 1998 and Mormann et al 2000.

In analogy to the directional coherence in Eq. 36, the “periodogram-based *phase* coherence” is:

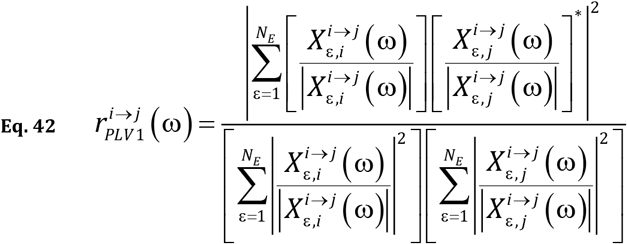

which simplifies to:

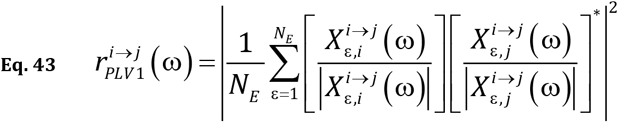

Note that these equations are based on the phases of the directional Fourier transforms in Eq. 26, which provide high frequency resolution. In this case, no use is made of the instantaneous phase. A similar form of equation, for the nondirectional case, appears in Lowet et al 2016, Equation 15 therein, and in Pascual-Marqui 2007, Equation 54 therein.

The next two definitions are based on the coherence between the directional instantaneous phases given by Eq. 34 and Eq. 35:

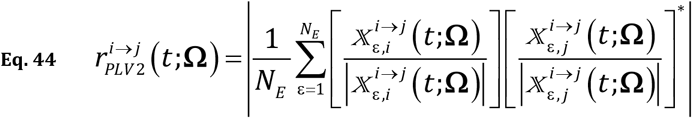

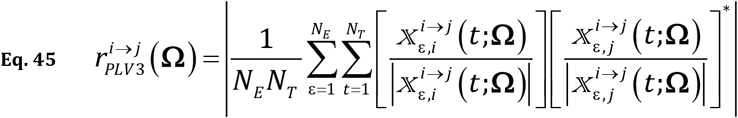

where Ω denotes the frequency band settings of the band-pass filter used in the algorithm “Alg2-Step# 1”.

The phase coherence in Eq. 44 is a function of time, while Eq. 45 corresponds to the average over time.

Directional cross-frequency versions for frequency pairs (ω_1_,ω_2_) and for band pairs (**Ω**_1_,**Ω**_2_) are defined as:

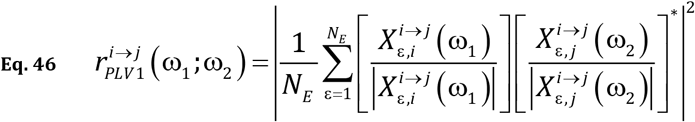

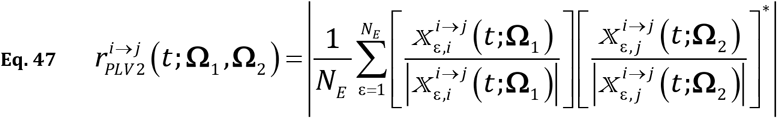

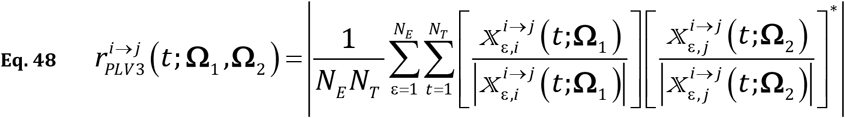

The special appeal of the directional PLV in Eq. 43 is its frequency resolution, which is much higher than that obtained by band pass filtering just prior to the discrete Hilbert transform, as required by the PLV in Eq. 44 and Eq. 45. However, the special appeal of the directional PLV in Eq. 44 is its high temporal resolution.

We are aware of one previous attempt to estimate the direction of information flow for phase-phase coupling, namely, the directed phase lag index (dPLI), by Stam and van Straaten (2012). It is based on a false premise: the sign of the phase difference (positive or negative) indicates the direction of flow. It suffices to change the sign of one signal to reverse the flow, which renders the dPLI meaningless. For instance, bipolar intracranial signals are determined up to a factor of ±1, rendering the dPLI meaningless between two signals each consisting of a local bipolar cortical electric potential difference. This fact was proven in Pascual-Marqui et al 2018.

The new directional measures of phase-phase coupling reported in this work are invariant to the sign of the signals.

Weighted directional PLVs for Eq. 43, Eq. 44, and Eq. 45 are:

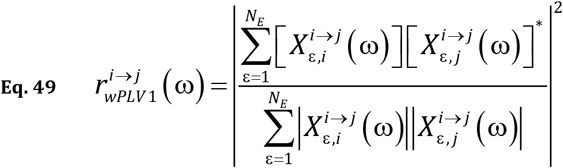

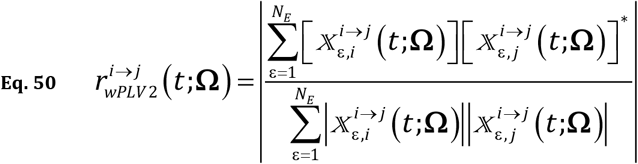

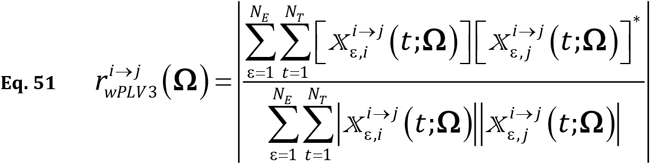

which are based on the work of Kovach 2017. Note that they bypass the calculation of phase, which is undefined for negligible amplitude.

### Directional phase-amplitude coupling (PAC) using linear causal filtering

There are many non-directional measures of phase-amplitude coupling (PAC) between two distinct signals. Taking an example from electrophysiology, and starting from two distinct signals, all the published measures quantify in different ways the non-directional association between two derived signals:

1. the envelope (instantaneous amplitude) of one signal that has been high-frequency band-pass filtered e.g. gamma 35 Hz to 55 Hz, and
2. the instantaneous phase of the other signal that has been low-frequency band-pass filtered e.g. theta 5 Hz to 7 Hz.

See e.g. Penny et al 2008 and Hülsemann et al 2019 for reviews and comparison of methods.

*The methods proposed here do not apply to phase-amplitude coupling of one signal. They apply to coupling between the phase of one signal, with the envelope of a distinctly different signal.*

To quantify the directional causal influence of the phase of the i-th signal on the amplitude of the j-th signal, one can use any non-directional measure of association between the directional phase and amplitude:

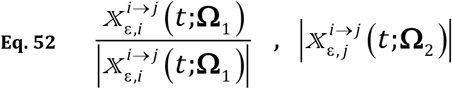

To quantify the directional causal influence of the amplitude of the j-th signal on the phase of the i-th signal, one can use any non-directional measure of association between the directional phase and amplitude:

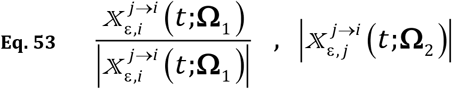

The quantities in Eq. 52 and Eq. 53 are defined in Eq. 32, Eq. 33, Eq. 34, and Eq. 35.

For instance, using the phase-amplitude coupling measure of Özkurt and Schnitzler (2011), the two distinct directional versions:

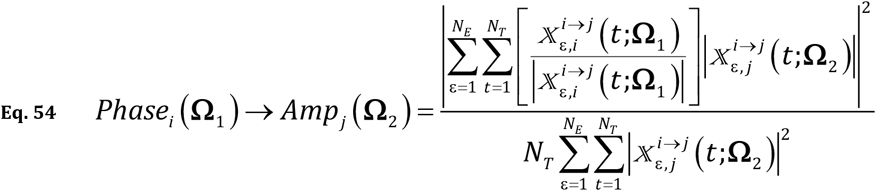

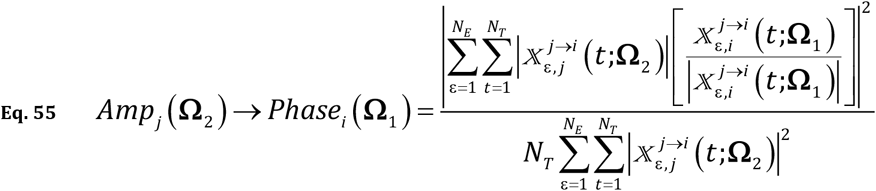

Eq. 54 quantifies the causal influence of the phase of the i-th signal on the envelope of the j-th signal. Eq. 55 quantifies the causal influence of the envelope of the j-th signal on the phase of the i-th signal. These measures can be used to distinguish which event is the cause of the coupling: for instance, does the phase of theta trigger gamma bursts, or are gamma bursts facilitating theta oscillations?

### Directional envelope-envelope coupling using linear causal filtering

The non-directional measure of envelope-envelope coupling between two signals is also known as “amplitude-amplitude coupling”. Some basic early literature on this non-causal, non-directional measure of association are the papers by Bruns et al 2000, and Bruns and Eckhorn 2004.

Taking an example from electrophysiology, starting from two distinct signals, this coupling quantifies the association between two derived real-valued signals:

1. the envelope (instantaneous amplitude) of one signal that has been band-pass filtered e.g. gamma 35 Hz to 55 Hz, and
2. the envelope (instantaneous amplitude) of the other signal that has been band-pass filtered e.g. theta 5 Hz to 7 Hz.

A simple measure of association is the simple cross-correlation between the two envelope signals, taking the value at the lag for which it is maximum, as described in Bruns and Eckhorn 2004.

Based on this measure, the directional envelope-envelope coupling is:

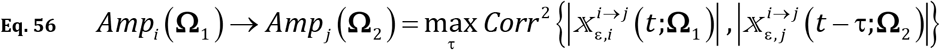

where *Corr*^2^ {●, ●} is the squared correlation coefficient between the two random variables, sampled over all admissible “t” and “ε”. The quantities in Eq. 56 are defined in Eq. 32 and Eq. 33.

For completeness, the squared correlation between *“y”* and “*z*”, for a total sample size *“N”,* is:

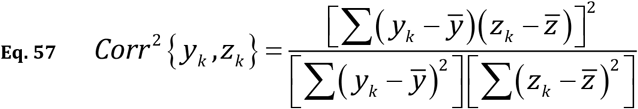

with:

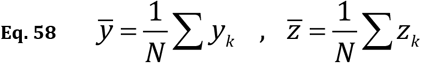

Eq. 56 quantifies the causal influence of the Ω_1_ -envelope of the i-th signal, on the Ω_2_ -envelope of the j-th signal.

The causal influence of the Ω_2_ -envelope of the j-th signal, on the Ω_1_ -envelope of the i-th signal can be quantified as:

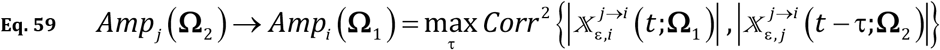

## 9. Discussion

The practical validity of the linear causal filter hinges on a set of very restrictive assumptions:

Assumption #1. Wide sense-stationarity.
Assumption #2. There are no other sources of influence other than those expressed by the signals in the multivariate autoregressive model.
Assumption #3. No measurement noise.

Actually, all Granger causality analyses hinge on these assumptions.

In particular, the measurement noise problem (Assumption #3) has been largely overlooked in the time series analysis literature. It seems to have been confused with a conventional simple “model plus noise” problem, where parameter estimation is typically unbiased.

Many popular “model plus noise” problems produce non-biased estimators using least squares techniques. This gives confidence in the estimators if the sample size if large enough.

However, for autoregressive models, least squares parameter estimation yields biased parameters. Other estimation techniques need to be used to obtain non-biased estimators (see e.g. Pascual-Marqui et al 2021, and references therein). Simple measurement noise can create false Granger causality, when using estimation technique that do not correctly account for measurement noise. The reason for this problem lies in the distinct nature of the innovations which determine the dynamics. If measurement noise is not explicitly modeled as such, it is confused with the innovations and incorrectly drives the dynamics, yielding false coupling coefficients.

The linear causal filter proposed here is parametric, in the sense that it relies on the multivariate autoregressive model and its coefficients. However, the same linear causal filter can be defined by using nonparametric minimum-phase spectral factorization, as developed by Dhamala et al (2008a, 2008b).

A generalization of the linear causal filter is straightforward for the case when there are three multivariate time series **x***(t*) ∈ ℝ^*p*×1^, **y***(t*) ∈ ℝ^*r*×1^, and **z**(*t*) ∈ ℝ^*s*×1^, where the causal effects of **z***(t*) are accounted for and partialled out, and where the causal coupling from **y**(*t*) to **x**(*t*) are severed, leaving only the two modified time series with only causal flow from “**x** to **y**”.

This multivariate generalization might be of interest in the analysis of brain activity signals. For instance, **x**, **y**, and **z** might represent activities from three different resting state networks (each one composed of several signals from different cortical areas). The linear causal filter will allow the study of the causal influence that one network has on another network, after accounting for and partialling out the effects of a third network. In this case, use must be made of multivariate non-directional measures of association for coherence and for cross-frequency couplings (e.g. phase-phase, phaseamplitude, amplitude-amplitude, etc), which can now be decomposed into their directional causal components.

Another possible generalization of the linear causal filter is its application to time varying autoregressive models, which can be implemented using e.g. the short time Fourier transform.

## 10. Appendix A: discrete Fourier transform (DFT)

The DFT for a time series *h*(*t*), for *t* = 0…(*N_T_* − 1), is defined as:

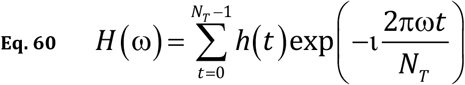

for ω = 0…(*N_T_* − 1), with ω = 0 corresponding to the DC component, ω = *N_T_*/2 corresponding to the Nyquist frequency, and with 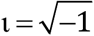. The DFT of a vector or of a matrix is calculated for each element. In practice, Eq. 60 is implemented by specialized very fast algorithms (see e.g. Cooley and Tukey 1965).

Let *S_R_* denote the actual sampling rate in Hz, i.e. in number of time samples per second. Then discrete frequency ω = 0…(*N_T_* − 1) corresponds to actual frequency *“f”* in Hz:

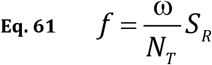

The DFT of a sequence of autoregressive coefficients *a*(*k*) of order *“q”* is a particular case which simplifies to:

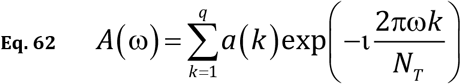

For real-valued time series, the periodogram estimator for the spectral density is:

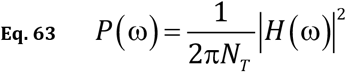

The tapered version of the DFT is:

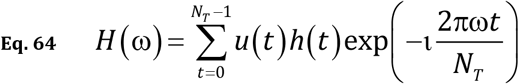

where *u*(*t*) for *t* = 0…(*N_T_* − 1) is the taper. A review on window functions (or tapers) *u*(*t*) can be found in Harris 1978. See also Brillinger 2001, and Oppenheim and Schafer 2014.

When using the tapered DFT, the periodogram estimator for the spectral density is:

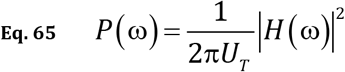

with:

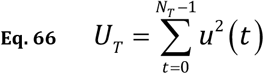

Note that when several epochs are available, the periodogram then corresponds to the average of Eq. 63 or Eq. 65 over epochs.

## 11. Appendix B: the inverse discrete Fourier transform (invDFT)

The invDFT of complex-valued *H*(ω), for ro = 0…(*N_T_* − 1), is defined as:

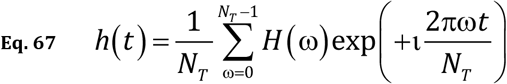

for *t* = 0…(*N_T_* − 1). In general, *h*(*t*) obtained from Eq. 67 can be complex-valued.

A real-valued time series *h*(*t*) is obtained when *H*(ω) is given for ro 0…(*N_T_*/2), and when the values at frequencies higher than the Nyquist frequency satisfy:

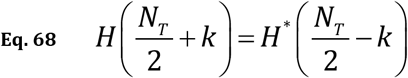

for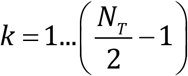.

If the invDFT is a zero mean real-valued time series, then follow these steps:

Step #1: Set:

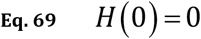

which corresponds to a zero mean time series (i.e. DC level equal to zero).

Step #2: Apply Eq. 68.

Step #3: Apply general definition of the invDFT in Eq. 67.

Step #4: End.

## 12. Appendix C: discrete Hilbert transform and the analytic signal

Let *x* (*t*) be a real valued signal. Let *y* (*t*) denote its Hilbert transform, defined as:

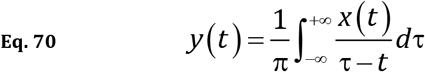

where the principle value of the integral is used.

The complex-valued analytic signal of *x* (*t*) is defined as:

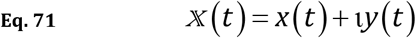

where 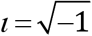.

In practice, for *x* (*t*) sampled at discrete time instants denoted *t* = 1…*N_T_*, a numerical algorithm for computing the analytic signal follows, based on Marple 1999.

**Alg3-Step# 1**

Compute the discrete Fourier transform of *x*(*t*), denoted *X*(ω) (see “Appendix A”).

**Alg3-Step# 2**

Define:

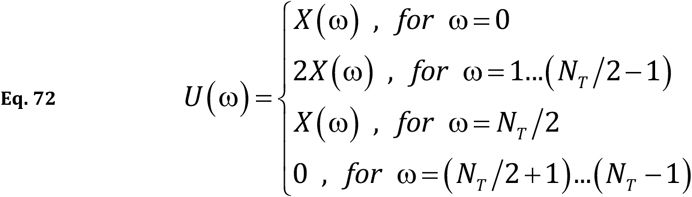

**Alg3-Step# 3**

The inverse discrete Fourier transform (see “Appendix B”) of *U*(ω) yields the complex-valued analytic signal 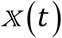 in Eq. 71 for *t* = 0…(*N_T_* − 1) .Note that the real part of 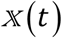 is the signal *x*(*t*), and the imaginary part *y*(*t*) is the discrete Hilbert transform.

**Alg3-Step# 4**

End.

The envelope, i.e. the instantaneous amplitude, is defined as:

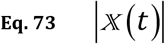

The complex valued instantaneous phase is defined as:

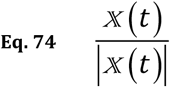

## 13. Acknowledgements

We greatly appreciate a detailed review of this manuscript by Daniele Marinazzo and Davide Nuzzi, which helped with its improvement, and importantly, revealed an error in the specification of the Fourier transform of the autoregressive coefficients, which has now been corrected.

## Graphical summary of the idea behind “Linear Causal Filtering”

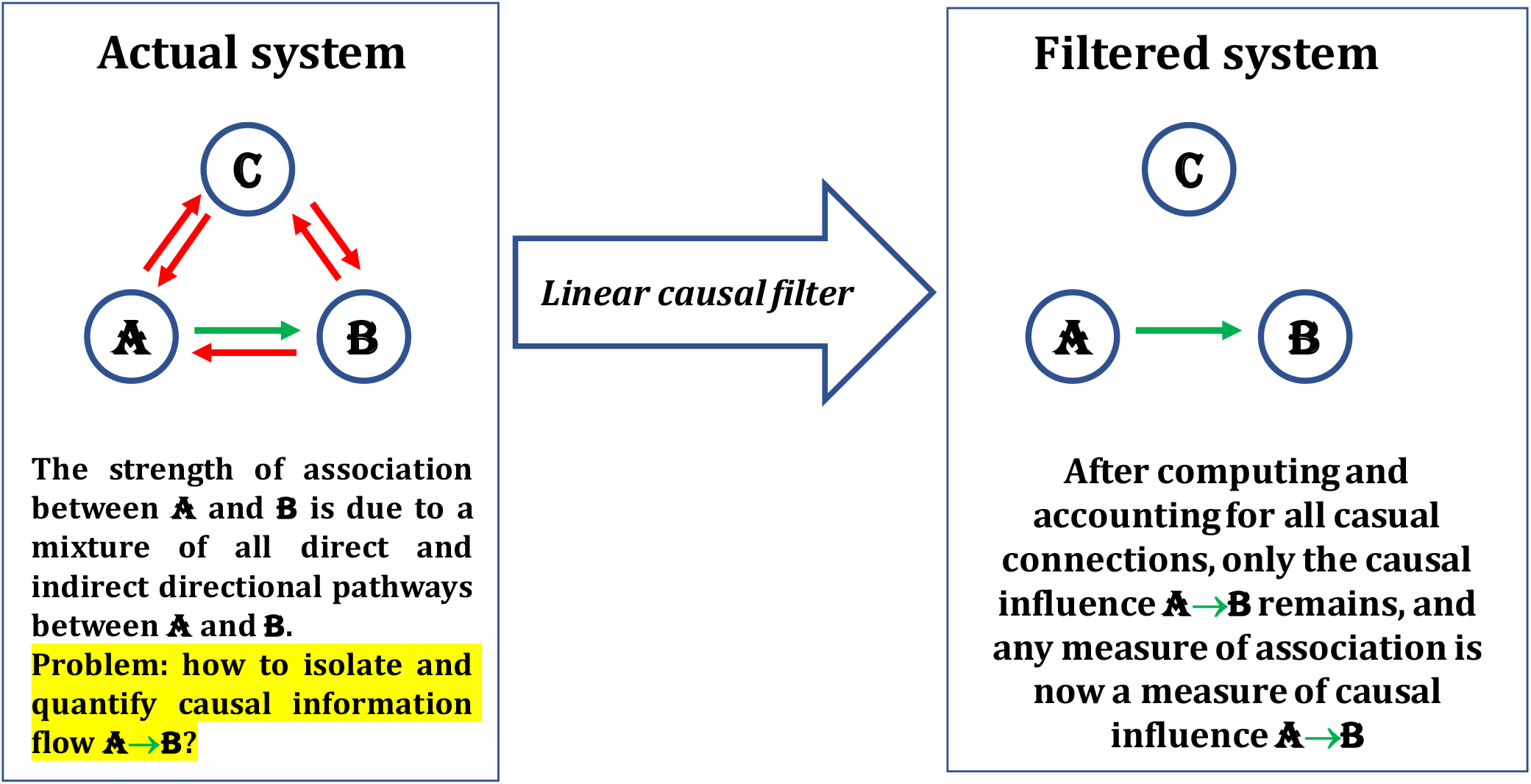

## Notes

### Competing Interest Statement

The authors have declared no competing interest.

